# Combining Gene Ontology with Deep Neural Networks to Enhance the Clustering of Single Cell RNA-Seq Data

**DOI:** 10.1101/437020

**Authors:** Jiajie Peng, Xiaoyu Wang, Xuequn Shang

## Abstract

**Background:** Single cell RNA sequencing (scRNA-seq) is applied to assay the individual transcriptomes of large numbers of cells. The gene expression at single-cell level provides an opportunity for better understanding of cell function and new discoveries in biomedical areas. To ensure that the single-cell based gene expression data are interpreted appropriately, it is crucial to develop new computational methods.

**Results:** In this article, we try to construct the structure of neural networks based on the prior knowledge of Gene Ontology (GO). By integrating GO with both unsupervised and supervised models, two novel methods are proposed, named GOAE (Gene Ontology AutoEncoder) and GONN (Gene Ontology Neural Network) respectively, for clustering of scRNA-seq data.

**Conclusions:** The evaluation results show that the proposed models outperform some state-of-the-art approaches. Furthermore, incorporating with GO, we provide an opportunity to interpret the underlying biological mechanism behind the neural network-based model.

## Introduction

In the past decade, transcriptome studies have benefited from next-generation sequencing (NGS) based RNA expression profiling (RNA-seq) [1]. However, the resulting expression value based on bulk RNA-seq is an average of its expression levels across a large population of input cells [2]. Such bulk expression profiles are insufficient to provide insight into the stochastic nature of gene expression [3]. Therefore, bulk measures of gene expression may not help researchers to understand the distinct function and role of different cells [2]. To address the problem, single cell RNA-seq (scRNA-seq) is applied to assay the individual transcriptomes of large numbers of cells [4]. The gene expression at single-cell level provides an opportunity for better understanding of cell function and new discoveries in biomedical areas [5, 6].

ScRNA-seq data analysis poses several new computational challenges. To ensure that the single-cell based gene expression data are interpreted appropriately, it is crucial to develop new computational methods. One of the most important applications of scRNA-seq is to group cells and identify new cell types. The major computational challenge in this application is clustering cells based on the gene expression at single-gene level. Clustering based on scRNA-seq data may help us understand underlying cellular mechanisms, which can promote the discovery of new markers on specific types of cells [7], and identification of tumor subtypes [8], etc.

In the clustering problem, cells are partitioned into different cell types based on their global transcriptome profiles. Each cell type has a significantly distinctive expression signature from the others. Since the expression values are always with high dimensionality and noise from the sequencing result, dimensionality reduction is usually performed before clustering. Till now, several methods have been proposed to eliminate the influence of noise and reduce the dimension. The existing methods could be loosely grouped into two categories, unsupervised method and supervised method.

In the unsupervised category, the main idea is to perform dimensionality reduction before clustering. The simplest method is based on the principal component analysis(PCA) [9]. As one of the most popular methods for dimensionality reduction, PCA has been studied extensively for clustering single cells [10–13]. Assuming that the data is normally distributed, PCA uses an orthogonal transformation to convert a set of observations of possibly correlated variables into a set of values of linearly uncorrelated variables, which are called principal components. However, for scRNA-seq datasets, they are not exactly linearly separable. T-distributed stochastic neighbor embedding (t-SNE) [14] is a nonlinear dimensionality reduction technique, which is also used for clustering single cells recently [12, 13]. Based on the Gaussian kernel, t-SNE converts high dimension data to low dimension space. But, it usually maps multidimensional data to two or more dimensions suitable for human observation. Hence it always accompanies with dimension restriction. Besides, similar with PCA, t-SNE also does not consider the drop out events of scRNA-seq data. To consider the specificity of scRNA-seq data, ZIFA [15] uses zero-inflated factors to deal with the drop out events in scRNA-seq data. Assuming that drop out events may lead to zero counts, ZIFA models these counts exactly zero rather than close to zero in the dataset. The evaluation test shows that ZIFA performs better than PCA and t-SNE on some datasets. But the hypothesis of ZIFA is that zero is inflated as Gauss distribution, and the transformation between descending dimension and original data is linear. Given the expression profiles of single cells, SNN-cliq computes the similarity between cells by using the concept of shared nearest neighbor (SNN), and implements clustering algorithm based on graph theory [16]. By combining multiple clustering methods, SC3 performs a consensus clustering which includes spectral transformation, k-means algorithm, and complete link approach to achieve high accuracy and robustness [17]. However, SC3 and SNN-cliq cannot build a relationship between data representation and quantity and property of cell types. Integrating PCA and hierarchical clustering, pcaReduce tries to improve the original PCA method by finding a connection between the PCA-based representations and the number of resolvable cell types. Meanwhile, denoising autoencoder (DAE) [12] is used to reconstruct the data from high dimensions to low dimension space.

Motivated by the success of neural networks in other areas, Lin et al. develop a supervised method to generate the low dimensional representation of scRNA-seq data based on neural networks (NN) [12]. NN model combines the neural network with the protein-protein interaction (PPI) network to classify a number of cells. Given cells with know cell types, this model can be trained as a supervised model. After that, the hidden layer of the trained neural networks is used for generating the low dimensional representation of scRNA-seq data. The experimental test shows that this supervised method performs better than most of existing unsupervised models.

Although many attempts have been made to cluster single cells based on the global transcriptome profiles, most of them only consider the transcriptome profiles neglecting the prior biological knowledge. This large limits the performance of state-of-art systems. Inspired by the success of neural network in modeling the hierarchical structure and function of a cell [18], we ask whether combining the rich prior biological knowledge in gene ontology (GO) with neural networks could enhance the clustering of cells based on their global transcriptome profiles. Gene Ontology (GO) [19], which has been widely used in many areas [20–24], provides a popular vocabulary system for systematically describing the attributes of genes and other biological entities. As one of the most popular bioinformatics sources, it contains reliable and easy-interpreted prior biological knowledge. In this article, we try to construct the structure of neural networks based on the prior knowledge of GO. By integrating GO with both supervised and unsupervised models, two novel methods are proposed, named GOAE (Gene Ontology AutoEncoder) and GONN (Gene Ontology Neural Network) respectively, for clustering of scRNA-seq data. The major contributions of this article are as follows:

- To better dimensionality reduction of scRNA-seq data, we propose a novel neural work structure considering the prior knowledge in GO.
- We propose two novel models, named GOAE and GONN, to enhance cluster cells based on their transcriptome profiles.
- The evaluation results show that the proposed models outperform some stateof-the-art approaches.
- Incorporating with GO, we provide an opportunity to interpret the underlying biological mechanism behind the neural network-based model.

## Methods

We propose a novel model to obtain the low dimensional representation of scRNA seq data by combining the Gene Ontology and neural network model. We use the terms in GO to replace the neuron in the neural network and convert the fully connected neural network as partial-connected. Based on this idea, we propose two novel methods: an unsupervised method based on an autoencoder model and a supervised method based on a traditional neural network model. The basic idea of our models is to perform the dimensionality reduction by training neural network (or autoencoder) model and extract the latent layer as low dimensional representation. This section consists of following components. First, we will introduce how to select significant GO terms from the whole GO structure. Second, we combine GO with an autoencoder to build an unsupervised model for dimensionality reduction, named GOAE. Third, we combine GO with a neural network to build a supervised model for dimensionality reduction, named GONN. Finally, the low dimensional representation is used for clustering of cells based on a clustering method.

**Table 1.** Number of Gene Ontology terms at different layers.

| layer number | 0 | 1 | 2 | 3 | 4 | 5 | 6 | 7 | 8 | 9 | 10 | 11 |
| --- | --- | --- | --- | --- | --- | --- | --- | --- | --- | --- | --- | --- |
| biology process | 1 | 24 | 151 | **1010** | 2662 | 3934 | 3443 | 2167 | 784 | 305 | 98 | 15 |
| molecular function | 1 | 17 | 111 | **476** | 894 | 1397 | 887 | 481 | 196 | 63 | 23 | 4 |

### Selection of significant GO terms

Gene Ontology (GO) is a popular vocabulary system for systematically describing the attributes of gene and gene product. Each GO term could annotate a set of genes. GO consists of three different categories, which are biology process, molecular function and cellular component. GO is structured as a directed acyclic graph. Each term has defined relations with other terms in the same or various categories. In this step, we choose GO terms that are used in the following model. We only use terms in the biological process and molecular function category since these terms might be more functional related. In GO, a parent term annotates all the genes annotated by its descendants. The main steps of selecting GO terms used in the following steps are as follows.

First, we select all the GO terms at the third layer. Evaluation test shows that GO terms at the third layer can achieve the best performance. The number of GO terms at different levels are shown in Table 1. These 1543 GO terms at the third level are the candidate terms that connect with the input layer in the neural network.

Second, we remove redundancy terms from the candidate terms obtained from last step. The annotated genes of different terms may have overlap. Therefore, we remove the redundancy terms to decrease the information redundancy and the parameters in the following neural network-based model.

Specifically, let *GO*_*i*_: {*gene*_1_, *gene*_2_, …*gene*_*n*_} be a GO term, named *GO*_*i*_, annotating a set of annotation genes *gene*_1_, *gene*_2_, …*gene*_*n*_. The unique score *S*_*u*_ of two GO terms are defined as follows:

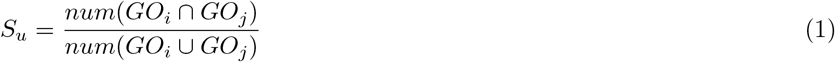

If the unique score *S*_*u*_ of two GO terms is larger than 0.5, the two GO terms are considered as not unique. Then, we will delete the GO term that has fewer annotation genes.

Third, we remove the terms annotating genes that have similar expression profiles in different cells. Different genes may have different expression level in different cells. We tend to select the genes that have different expression levels for clustering. Therefore, we select the terms annotating genes having diverse expression levels in different cells. The diversity of a GO terms could be measured by gene expression values. z-score-based method is used for normalization on gene dimension. Following this normalize operation, the expression values of each gene is normalized as a standard normal distribution. We define *std*_*j*_ as standard deviation of *gene*_*j*_. The diversity score *H*_*i*_ of a GO term *GO*_*i*_ is calculated as follows:

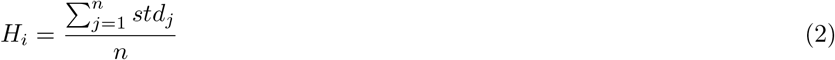

where *n* is the number of genes annotated by *GO*_*i*_. If the diversity score of *GO*_*i*_ is less than the given threshold (in this case 0.1), *GO*_*i*_ is considered as low diversity term. We then delete the low diversity GO terms.

After these three steps, we obtain a set of GO terms with low redundancy and high diversity.

### Architecture of unsupervised model (GOAE)

In the task of scRNA-seq data clustering, an unsupervised dimensionality reduction model is a key component. To perform the dimensionality reduction, we combine the Gene Ontology with autoencoder that have been widely used in other areas, like image processing, natural language processing.

To combine the GO with neural network, we add GO terms to the neural network as partial-connected neurons. The structure of this model is formulated from extensive prior knowledge of gene ontology. The architecture of GOAE is shown in figure 1.

The input layer are genes involved in the scRNA-seq datasets. In hidden layer 1, BP neurons and MF neurons are added based on the biological process and molecular function terms obtained from GO. As shown in Figure 1, BP and MF neurons are partially connected. Only genes annotated by the corresponding GO term are feeded to the GO term neuron.

GOAE consists of two components: encoder and decoder. From the input layer to hidden layer 2 are the encoder. The decoder part is exactly a mirror of the encoder part, which from hidden layer 2 to the output layer.

Let *x*_*i*_ be the output of the *i*th hidden layer. The forward propagation of the neural network is:

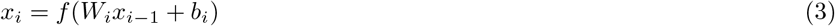

where *W*_*i*_ represents the weight matrix of the edge from *i* − 1 th layer to *i*th layer in the neural network, *b*_*i*_ is the bias of each *i*th hidden layer node, *f* (·) is the activation function. We choose tanh function in our case, which is:

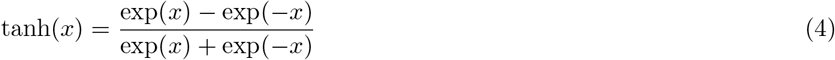

In this GOAE model, we use the mean square error as a loss function. Let *x*_0*j*_ be the input vector of sample *j*, and *x*_4*j*_ is the output vector. *n* represents the number of training sample. The loss function is defined as follows:

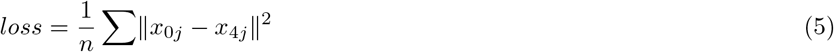

After several training epochs, the hidden layer 2 could be a low-dimension space of the input data.

Since the encoder and decoder are completely symmetric, both input layer and output layer are partial connection.

After training GOAE model, the hidden layer 2 could be used as the low-dimension representation of a cell. Then we can use a clustering method, (in our case, kmeans++), for the clustering of single cells.

**Figure 1.**
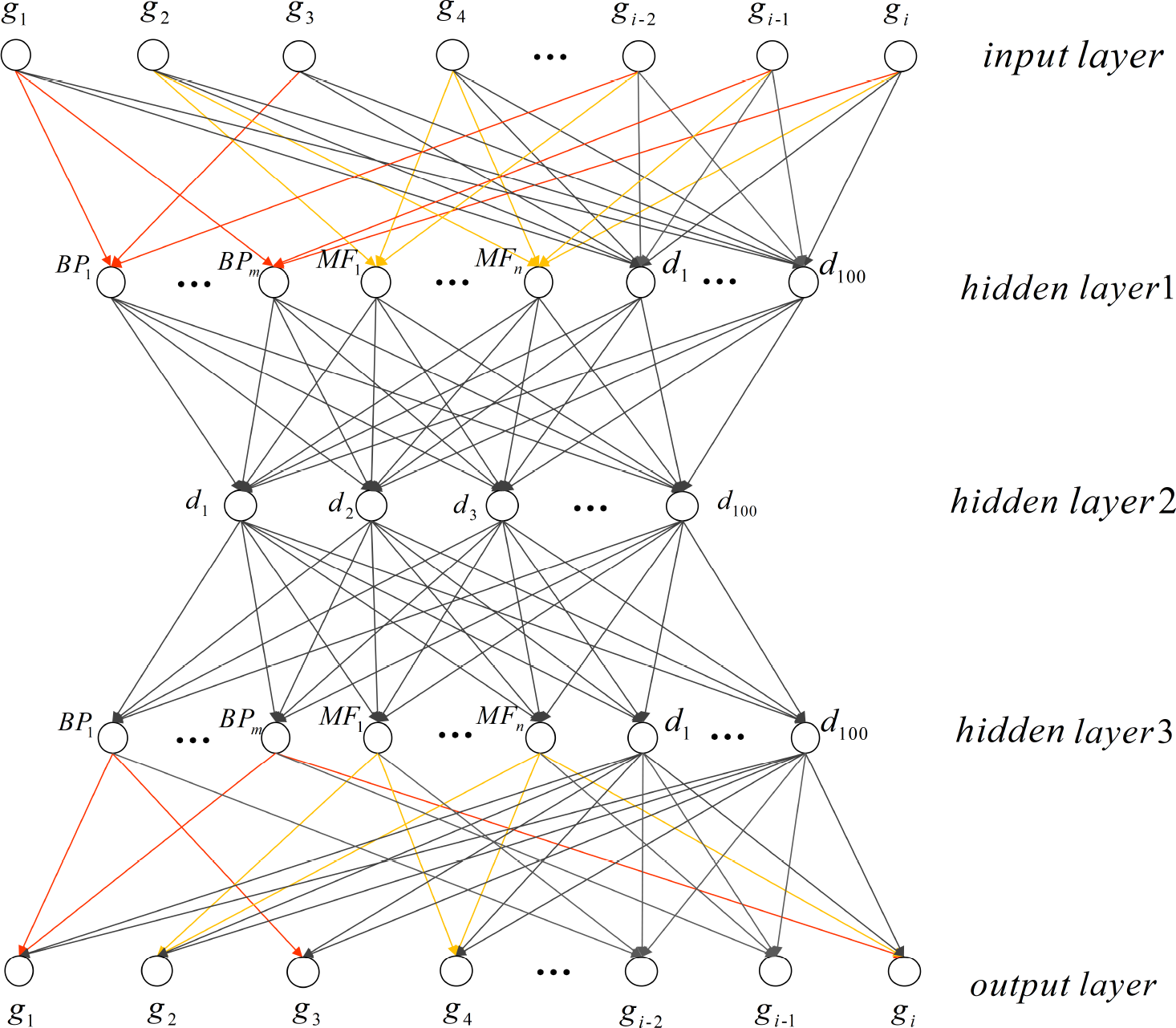
Gene Ontology Autoencoder (GOAE) model architecture. *BP*_*m*_ and *MF*_*n*_ are the GO-term neurons corresponding to biological process and molecular function category respectively. These neurons partially connected with the input layer. Node *d* is the full connected neurons. *g*_*i*_ represents the input gene in the given dataset.

### Architecture of supervised model (GONN)

A supervised dimensionality reduction model may also be needed in single cell clustering or retrieval [12]. Similar with the GOAE model, we replace the hidden layer1 neurons of the neural network with GO term nodes, which are partial-connected to the input layer neurons that represents the genes. In the GONN model, another hidden layer with 100 fully-connected neurons are added (see Figure 2). After the training phase, the hidden layer with 100 fully-connected neurons is considered as the low dimensional representation of the input.

At the output layer, softmax function is used for classification. Softmax function is defined as:

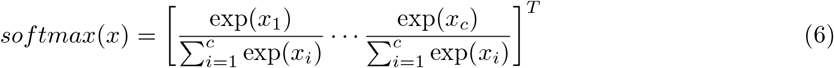

where *x* is the input vector of output layer and *c* is the number of all cell types. Based on softmax activation function, we can obtain the probability vector that a cell is classified into different cell types. Finally, we use top-1 method (the label which has the largest probability) to decide the cell type of a cell. In GONN, the loss is defined as:

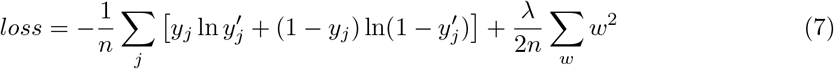

where *n* is the number of samples in the training dataset. The first part of Equation 7 is cross entropy. *y_j_* and 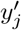 represent the desired output and the predicted output of sample *j* respectively. The second part is L2 regularization, where *λ* is the L2 regularization coefficient. *w* represents the training parameter vector. We combine cross entropy and L2 regularization to avoid overfitting and optimize parameters.

**Figure 2.**
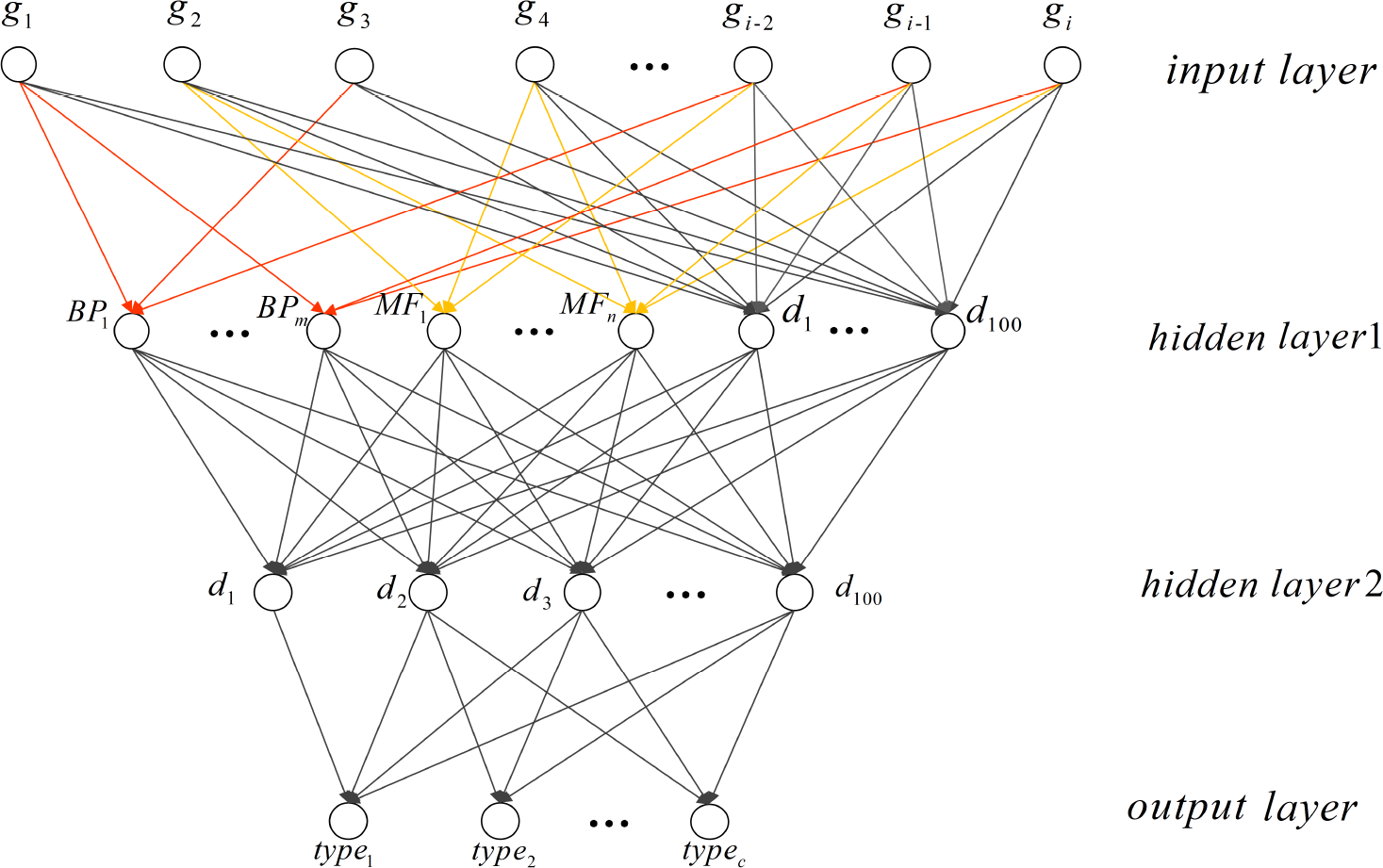
GONN model architectures. *type*_*c*_ represents different cell types as the label in the output layer. *BP*_*m*_, *MF*_*n*_, *d* and *g*_*i*_ are illustrated in Figure 1.

After training GONN models by known label cells, we extract the information of the last hidden layer(hidden layer2) as the low-dimension representation. Then we can use a clustering method, (in our case, kmeans++), for the clustering of single cells.

## Result

We test our models on two different scRNA-seq datasets. We compare our methods with two supervised methods (i.e. NN(ppi/tf) [12] and NN(dense)) and six unsupervised methods(i.e. PCA [9], t-SNE [14], ICA [25], pcaReduce [26], ZIFA [15], DAE [27]). For NN(dense) model, it has the same architecture as the two-layer GONN model but without partial connection between the input layer and hidden layer1. The NN(dense) model is used to test whether combining GO information can improve the supervised model. The DAE model is used to test whether the addition of GO information can improve the unsupervised neural network model. We also compare our model with other unsupervised methods. In all tests, we use *kmeas*++ for clustering based on different low-dimensional representations from different dimensionality reduction methods. The models are implemented using Python 3.6 and tensorflow 1.4.1 package.

### Data preparation

We test our models on two scRNA-seq datasets. One dataset is human scRNAseq data from [28]. In our experiment, 300 cells involving 11 cell types are used. The involved cell types are listed as follows: CRL-2338(epithelial), CRL2339(lymphoblastoid), BJ(fibroblast from human foreskin), GW(gestational 16, 21, 21+3 weeks from fetal cortex), HL60(myeloid from acute leukemia), iPS(pluripotent), K562(myeloid from chronic leukemia), Kera(foreskin keratinocyte) and NPC(neural progenitor cells). We remove the genes that have missing values in these cell types. 8686 genes are involved in the evaluation dataset. The other dataset is obtained from [12]. It integrate three mus musculus scRNA-seq datasets [11, 29, 30], which contains 402 cells involving 16 cell types. Similarly, after removing the genes with missing values, 9437 genes are included in the evaluation dataset. The gene ontology data is downloaded from *http://www.geneontology.org/*.

We set batch size as 64, epoch number as 100, learning rates as 1e-3 for GOAE model. We set the batch size as 64, epoch number as 200, learning rates as 0.2 for GONN model.

### Evaluation criteria

We use the adjusted rand index(ARI) [31] to compare the clustering results of single cells with true labels. ARI score can measure the similarity between two clustering results. It is defined as follows. Let *X* = {*X*_1_, …, *X*_*r*_} and *Y* = {*Y*_*1*_, …, *Y*_*s*_} be two different clustering results. *n*_*ij*_ represents the number of objects in common between *X*_*i*_ and *Y*_*j*_. Let *a*_*i*_ = ∑_*j*_ *n*_*ij*_ and *b*_*j*_ = ∑_*i*_ *n*_*ij*_, the ARI is defined as follow:

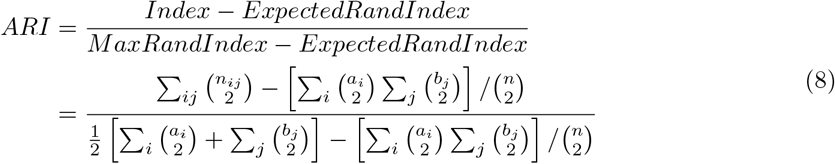

The scale of ARI score is between −1 and 1. The higher the ARI score is, the more similar two clustering results are.

Furthermore, normalized mutual information(NMI) [32] is also used for evaluation. NMI uses the concept of information entropy to compare different clustering results. NMI score is calculated as follows.

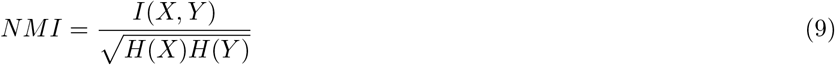

*H*(*X*) is the entropy of X, which is calculated as follows.

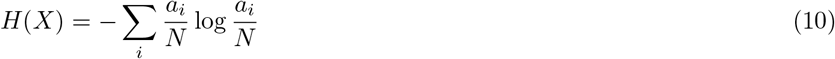

**Figure 3.**
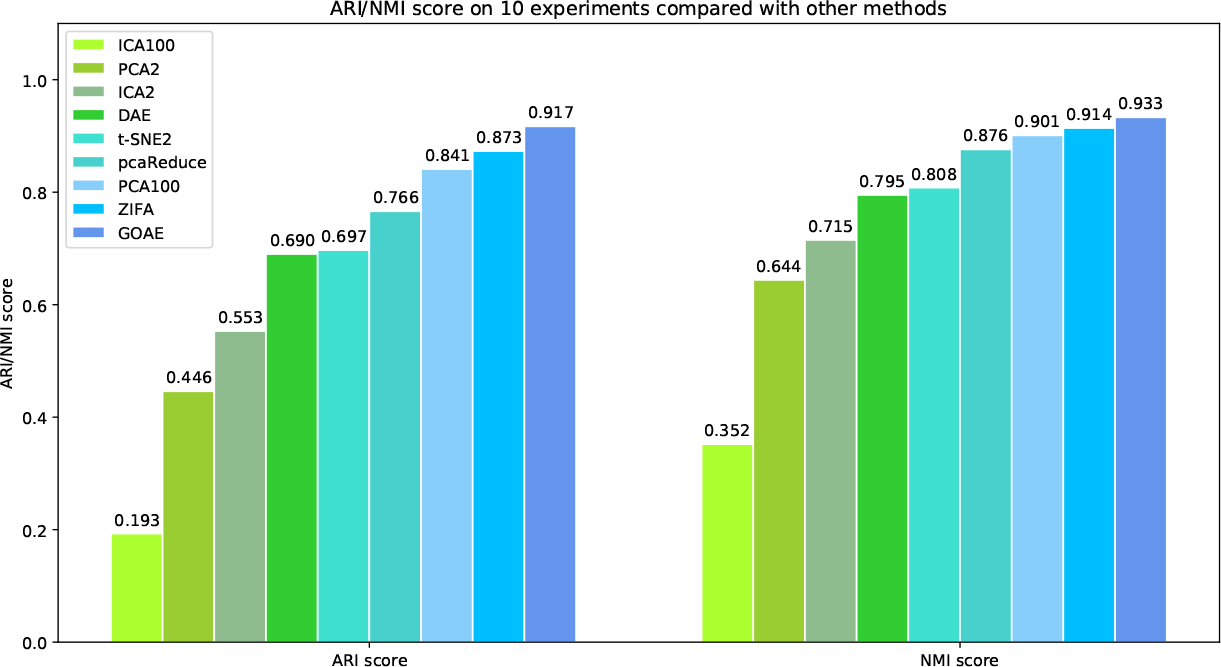
Average ARI and MNI scores on 10 experiments of different unsupervised methods on clustering 11 types of human cells. The number after methods means n components. i.e.PCA2 means using PCA method which we set 2 components.

*I*(*X*, *Y*) is the mutual information between *X* and *Y*, which is calculated as follows.

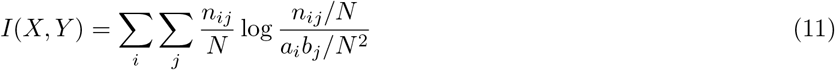

where *N* = ∑_*i*_ ∑_*j*_ *n*_*ij*_. NMI scores are between 0 and 1. The higher the NMI score is, the more similar two clustering results are. In the following evaluations, we run each experiment 10 times and calculate their average scores as final result.

### Performance evaluation on human scRNA-seq dataset

We test GOAE model (Figure 1) and GONN model (Figure 2) for clustering of human cells. 1174 GO terms satisfy the criteria described in 2.1 subsection. These terms are used in the GOAE and GONN model.

In the unsupervised test, all the unsupervised models are applied to the whole data set. All 11 types of cells are involved. Overall, GOAE performs the best among all tested methods. Similar with the experiment design in [12], several possible parameters (number of components) are tested for PCA and ICA method. We reduce the dimension of all data and using kmeans++ method to cluster all 11 cell types data. Figure 3 shows that GOAE perfects the best among all tested methods. The ARI and NMI score of GOAE are 0.917 and 0.933 respectively, while the scores of the runner-up method ZIFA are 0.873 and 0.914 respectively. The experiment result indicates that combining Gene Ontology and autoencoder can improve the performance of clustering of single cells.

For the supervised model, we compare GONN with the state-of-art method NN(ppi/tf) [12] and the original neural network model (NN). We apply the same experimental protocol used in [12]. The cell types not used in the training phase are used as the test set. There are 11 cell types involved in this data set. We randomly select 2, 4 and 6 cell types as the test set in the evaluation test.

**Table 2.**
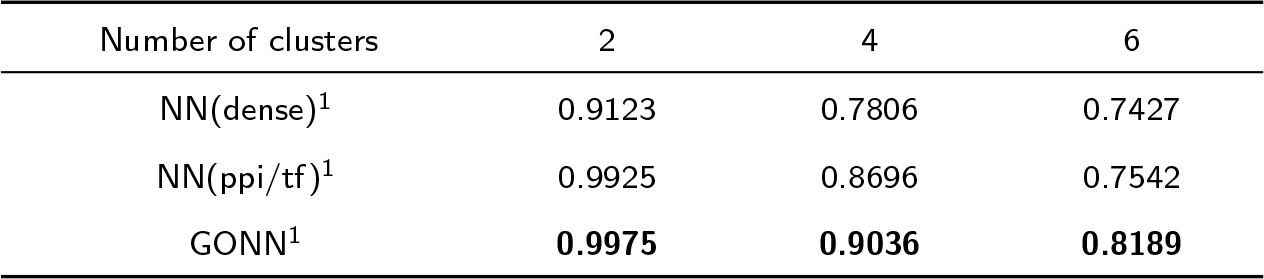
Average ARI scores of 10 experiments compared with other supervised model on human scRNA-seq dataset.For NN(dense) model, we set epoch number=200, learning rate=0.2. For NN(ppi/tf) model, the parameters are same as [12] epoch number=100, learning rate=0.1. For GONN model, we set epoch number=200, laerning rate=0.2.

**Table 3.**
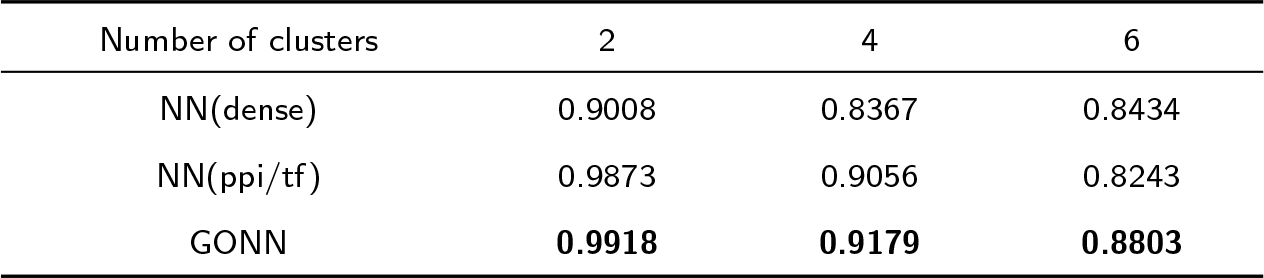
Average NMI scores of 10 experiments compared with other supervised model on human scRNA-seq dataset. See Table 2 for the hyper parameter selection of each model.

**Figure 4.**
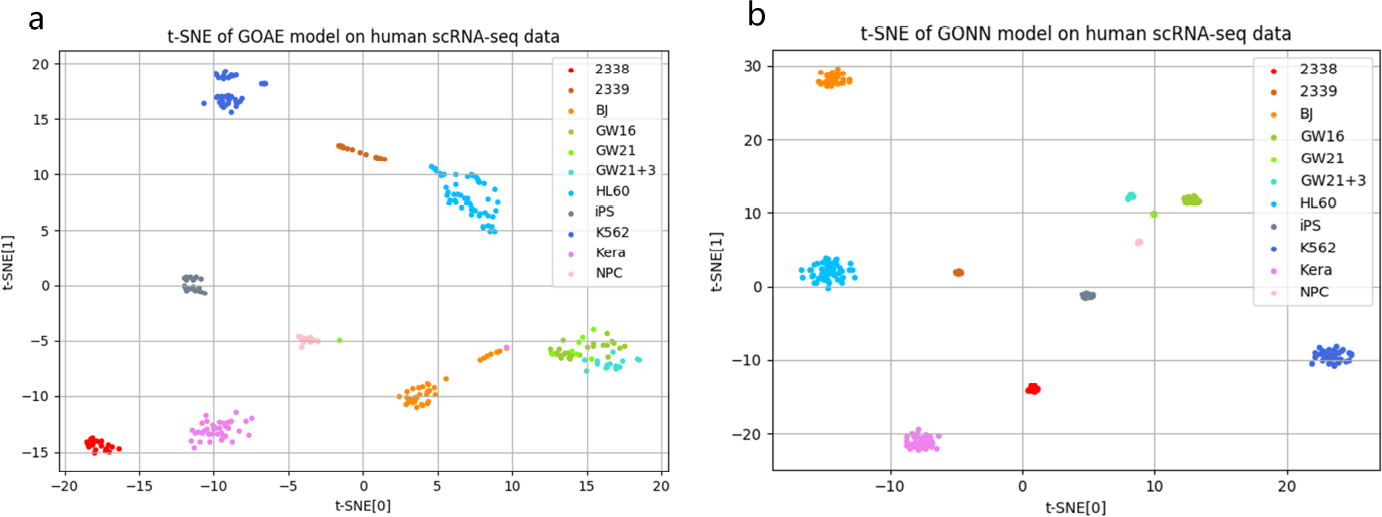
2D t-sne visualizations for our model on human scRNA-seq data. (a) is the dimension reduction result on GOAE model. (b) is the dimension reduction result on GONN model.

Overall, GONN method performs better than other methods (Table 2, 3). With the increase of number of cell types in the test set, the clustering task becomes more challenging. The result shows that GONN performs the best when the number of cell types equals to 2, 4 and 6. Furthermore, when the number of cell types is 6, the ARI score of GONN is 0.8189, which is significantly higher than the runner-up method (Table 2). Unsurprisingly, GONN method also achieves the highest NMI score. The NMI score of GONN is 0.8803 even when the number of cell types is 6, while the value of the second best method is 0.8434.

Figure 4 is the 2D visualization of low dimensional representation based on GONN and GOAE. We use t-SNE as the visualization tool. It is shown that the single cells are partitioned into different clusters based on GONN and GOAE, indicating that GONN and GOAE can learn a low dimensional representation for single cell data.

**Table 4.**
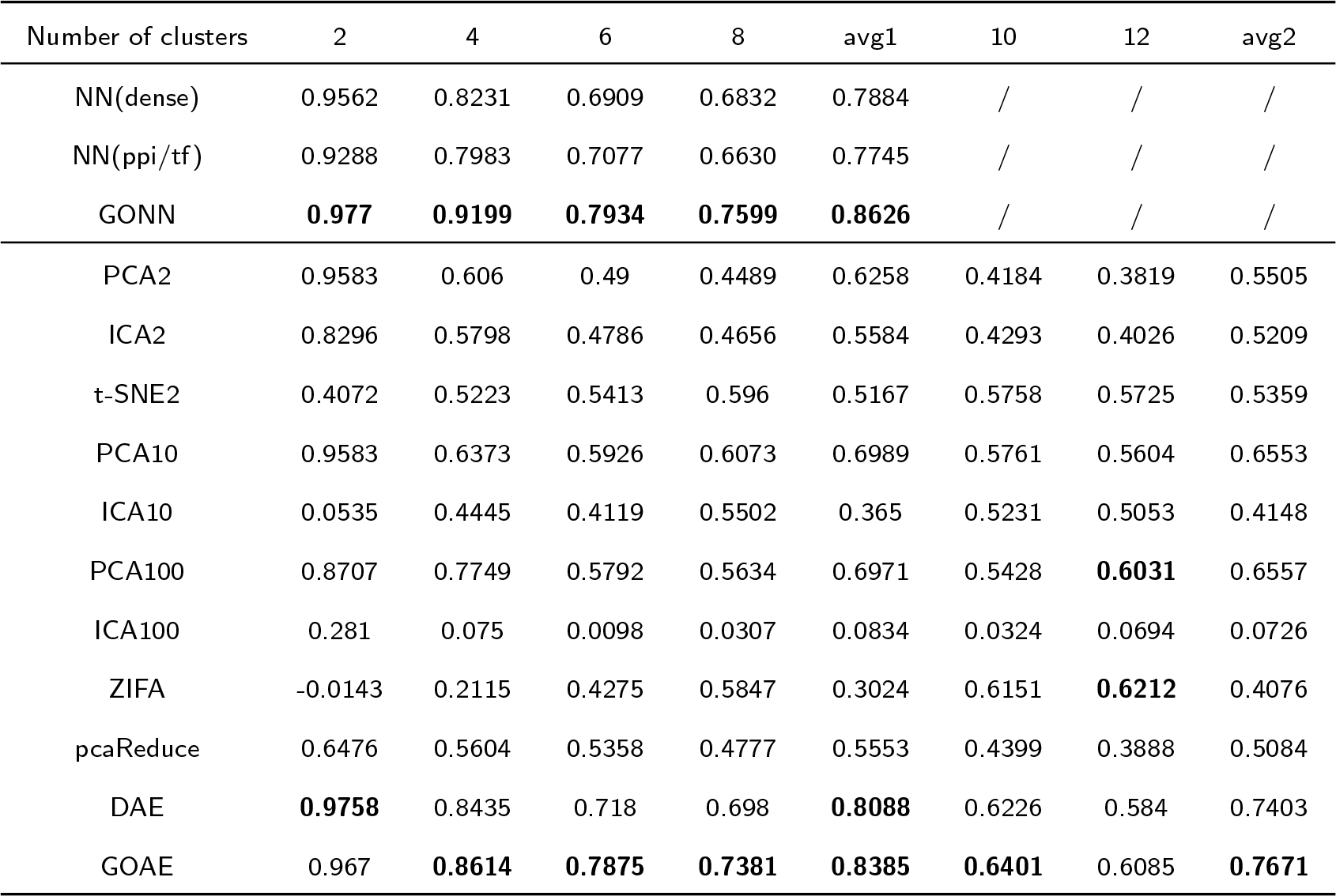
Average ARI score of 10 experiments compared with other methods on mus musculus dataset. The number after other unsupervised methods means n components. i.e.PCA2 means using PCA method which we set 2 components. For DAE model, we set epoch number as 200 and learning rate as 1e-3. For GOAE model, we set epoch number as 100 and learning rate as 1e-3. The parameters in other NN-based models are shown in Table 2. Avg1 is the average ARI score of the formal four cluster results, while avg2 ARI score is the average of all 2,4,6,8,10 and 12 cluster results. The highest values are shown in boldface.

| Number of clusters | 2 | 4 | 6 | 8 | avg1 | 10 | 12 | avg2 |
| --- | --- | --- | --- | --- | --- | --- | --- | --- |
| NN(dense) | 0.9562 | 0.8231 | 0.6909 | 0.6832 | 0.7884 | / | / | / |
| NN(ppi/tf) | 0.9288 | 0.7983 | 0.7077 | 0.6630 | 0.7745 | / | / | / |
| GONN | **0.977** | **0.9199** | **0.7934** | **0.7599** | **0.8626** | / | / | / |
| PCA2 | 0.9583 | 0.606 | 0.49 | 0.4489 | 0.6258 | 0.4184 | 0.3819 | 0.5505 |
| ICA2 | 0.8296 | 0.5798 | 0.4786 | 0.4656 | 0.5584 | 0.4293 | 0.4026 | 0.5209 |
| t-SNE2 | 0.4072 | 0.5223 | 0.5413 | 0.596 | 0.5167 | 0.5758 | 0.5725 | 0.5359 |
| PCA10 | 0.9583 | 0.6373 | 0.5926 | 0.6073 | 0.6989 | 0.5761 | 0.5604 | 0.6553 |
| ICA10 | 0.0535 | 0.4445 | 0.4119 | 0.5502 | 0.365 | 0.5231 | 0.5053 | 0.4148 |
| PCA100 | 0.8707 | 0.7749 | 0.5792 | 0.5634 | 0.6971 | 0.5428 | **0.6031** | 0.6557 |
| ICA100 | 0.281 | 0.075 | 0.0098 | 0.0307 | 0.0834 | 0.0324 | 0.0694 | 0.0726 |
| ZIFA | -0.0143 | 0.2115 | 0.4275 | 0.5847 | 0.3024 | 0.6151 | **0.6212** | 0.4076 |
| pcaReduce | 0.6476 | 0.5604 | 0.5358 | 0.4777 | 0.5553 | 0.4399 | 0.3888 | 0.5084 |
| DAE | **0.9758** | 0.8435 | 0.718 | 0.698 | **0.8088** | 0.6226 | 0.584 | 0.7403 |
| GOAE | 0.967 | **0.8614** | **0.7875** | **0.7381** | **0.8385** | **0.6401** | 0.6085 | **0.7671** |

### Performance evaluation on mus musculus dataset

Similar with evaluation test on human dataset, we also test these models on mus musculus dataset that contains 16 cell types. For unsupervised models, we randomly select 2, 4, 6, 8, 10 and 12 cell types as test sets. For supervised models, since sufficient training set is necessary, we only randomly select 2, 4, 6 and 8 cell types as test sets. The rest of data are used as the training set. For GOAE and GONN model, 854 GO terms satisfy the criteria described in 2.1 subsection.

As shown in Table 4, 5, for the unsupervised model, GOAE achieves the highest performance on datasets with different numbers of cell types. The average of ARI scores of GOAE on all datasets is 0.7671, which is around 0.03 higher than the runner-up method DAE. More details are shown in Table 4. The trend of NMI scores is similar to ARI scores. GOAE can achieve the highest NMI scores on datasets with different numbers of cell types. The complexity of the problem increases with the increase of the number of cell types. When the number of cell types is 8, the NMI score of GOAE is 0.8545 that is 0.04 higher than the runner-up method DAE. The evaluation test on mus musculus dataset indicates that combining gene ontology with neural network can improve the performance of single cell RNA-seq data clustering.

For the supervised model, GONN performs better than other compared methods. The ARI score decreases with the increase of the number of cell types involved in the test set. GONN can achieve a high ARI score (0.7599) even the number of cell types is 8, while the value of runner-up method is 0.6832. Similarly, GONN also achieve the highest NMI score in all tested methods. The average NMI score of different datasets is 0.9103, which is significantly higher than *NN* (*dense*) and *NN* (*ppi/tf*) method. The corresponding values of *NN* (*dense*) and *NN* (*ppi/tf*) are 0.8623 and 0.8496 respectively.

**Table 5.** Average NMI scores on 10 experiments compared with other methods on mus musculus dataset. The highest values are shown in blodface. See Table 4 for the more details.

| Number of clusters | 2 | 4 | 6 | 8 | avg1 | 10 | 12 | avg2 |
| --- | --- | --- | --- | --- | --- | --- | --- | --- |
| NN(dense) | 0.9348 | 0.8794 | 0.8179 | 0.8171 | 0.8623 | / | / | / |
| NN(ppi/tf) | 0.9083 | 0.8673 | 0.8119 | 0.811 | 0.8496 | / | / | / |
| GONN | **0.9688** | **0.9366** | **0.8721** | **0.8635** | **0.9103** | / | / | / |
| PCA2 | 0.9527 | 0.727 | 0.673 | 0.6756 | 0.7571 | 0.6567 | 0.6408 | 0.7210 |
| ICA2 | 0.8374 | 0.6966 | 0.6635 | 0.6828 | 0.7201 | 0.6632 | 0.6553 | 0.7261 |
| t-SNE2 | 0.4025 | 0.6101 | 0.6778 | 0.734 | 0.6061 | 0.7432 | 0.7531 | 0.6467 |
| PCA10 | 0.9527 | 0.7574 | 0.7349 | 0.758 | 0.8008 | 0.7425 | 0.7382 | 0.7835 |
| ICA10 | 0.1367 | 0.6224 | 0.6095 | 0.7196 | 0.5221 | 0.709 | 0.6965 | 0.5737 |
| PCA100 | 0.8656 | 0.8509 | 0.7675 | 0.7812 | 0.8163 | 0.7794 | 0.8215 | 0.8118 |
| ICA100 | 0.2186 | 0.1319 | 0.1173 | 0.142 | 0.15245 | 0.1539 | 0.249 | 0.1665 |
| ZIFA | 0.0548 | 0.3721 | 0.6267 | 0.7716 | 0.4563 | 0.8188 | **0.8271** | 0.5611 |
| pcaReduce | 0.6645 | 0.7115 | 0.6853 | 0.6649 | 0.6816 | 0.6499 | 0.6247 | 0.6689 |
| DAE | **0.9621** | 0.8877 | 0.8172 | 0.8194 | 0.8716 | 0.8047 | 0.7957 | 0.8478 |
| GOAE | 0.9544 | **0.9076** | **0.8693** | **0.8545** | **0.8965** | **0.8206** | 0.8018 | **0.8680** |

**Figure 5.**
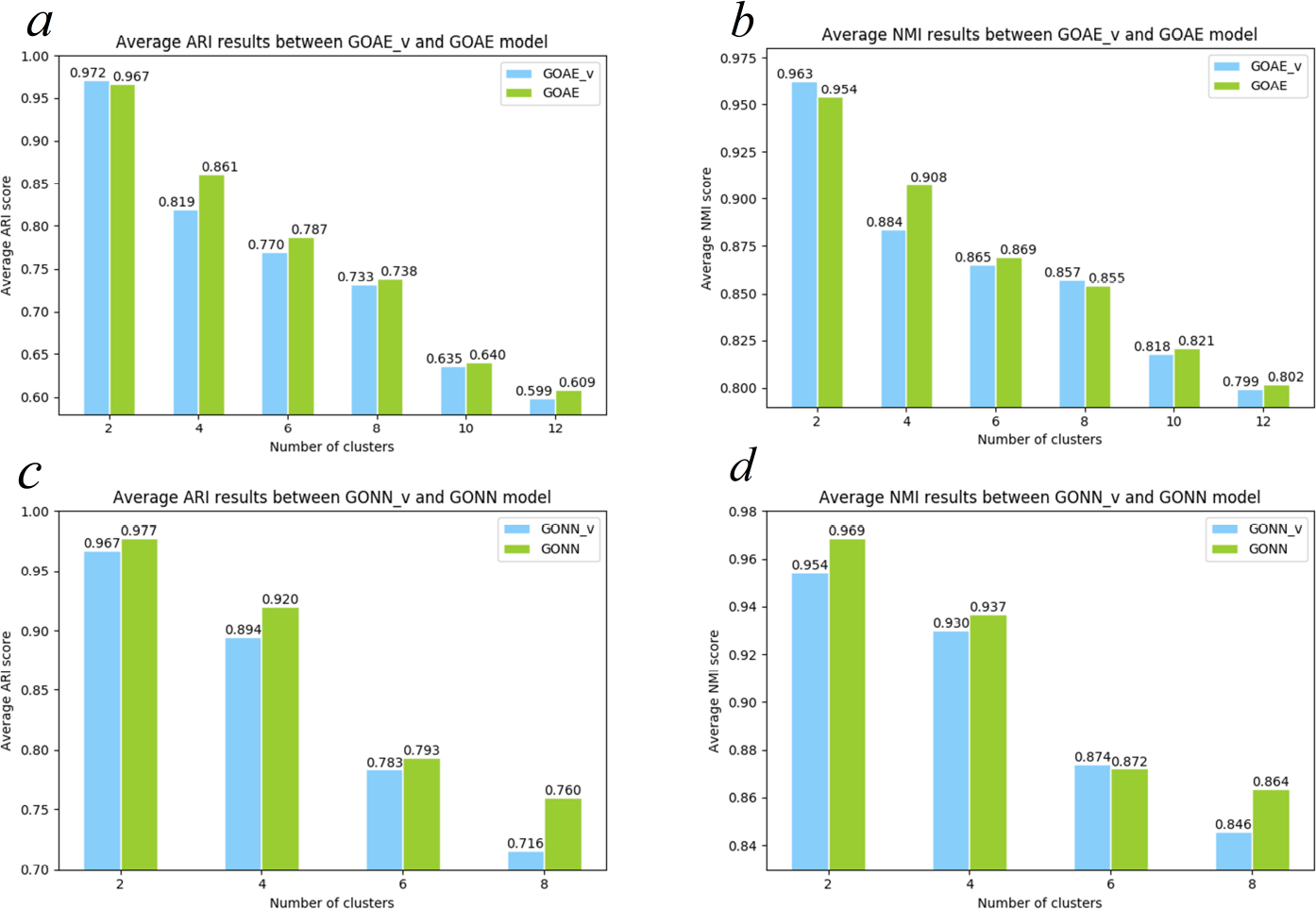
Performance evaluation by selecting different numbers of GO terms for GOAE and GONN model. (a, b) Average ARI and NMI results between *GOAE*_*v*_ model and *GOAE* model. *GOAE*_*v*_ represent GOAE model without selection of GO terms. (c, d) Average ARI and NMI results between *GONN*_*v*_ model and *GONN* model. *GONN*_*v*_ represent GONN model without selection of GO terms. GOAE models in (a) and (b) select different hyperparameters. Before GO terms selection, it selects epoch number=300, learning rate=1e-4. After selection, it selects epoch number=100, learning rate=1e-3. All GONN models in (c) and (d) all select epoch number=200 and learning rate=0.2. All these hyperparameters selection are achieve their best across a lots of hyperparameter groups.

**Table 6.** Highly ranked GO-term nodes for some cell types used for traing GOAE and GONN models

| Model | Cell type | GO term | GO function |
| --- | --- | --- | --- |
| GOAE | ES | GO:0043008 | ATP-dependent protein binding [33] |
|  | ES | GO:0032409 | Regulation of transporter activity [33] |
| GONN | ES | GO:0022417 | Protein maturation by protein folding [33] |
|  | ES | GO:0140101 | Catalytic activity, acting on a tRNA [33] |
|  | BMDC | GO:0099590 | Neurotransmitter receptor internalization [34] |
|  | BMDC | GO:0050881 | Musculoskeletal movement [34] |
|  | Zygote | GO:0001892 | Embryonic placenta development [35] |
|  | Early 2cell | GO:0032552 | Deoxyribonucleotide binding [36] |

### Effect of GO terms

One of the major contributions of our work is to add GO terms as neurons in the neural networks. To test whether the GO terms are selected appropriate, we re-run GONN and GOAE by varying the GO terms involved in the model. We use the mus musculus dataset on this test. As described in subsection 2.1, we remove the redundancy GO terms and GO terms with low diversity scores. In this test, we create *GONN*_*v*_ and *GOAE*_*v*_ where the redundancy and low-diversity GO terms are not removed. In *GONN*_*v*_ and *GOAE*_*v*_, 1486 GO terms are involved, while only 854 GO terms involved in GONN and GOAE. Figure 5(a) and (b) show that GONN is clearly better than *GONN*_*v*_, indicating that selecting appropriate GO terms contributes to the performance and this step has been appropriately designed. Similarly, Figure 5(c) and (d) show that GOAE is clearly better than *GOAE*_*v*_. Particularly, on the datasets with 8 and 10 cell types, the average ARI of GOAE are about 2-3% higher than *GOAE*_*v*_.

### Functional analysis on hidden layer nodes

For GOAE model, we train the model using samples of a certain cell type. Then, we could also obtain the top 10 highest GO-term nodes of the hidden layer. We select 8cell, 16cell, ES, earlyblast, and lateblast in this test, since training the GOAE model requires a sufficient amount of samples. For GONN model, we multiply the weight matrices *W*2 and *W*3 to represent the degree of importance between each cell type and the GO terms in the hidden layer 1. For each cell type, we selected the top-10 important GO terms for analysis. Table 6 shows some of the highly weighted GO-term nodes in the GOAE and GONN models. For example, regulation of transporter activity (GO:0032409) is mainly associated with ES(embryonic stem cell) [33], and embryonic placenta development (GO:0001892) is always relative with zygote cell [35].

## Conclusion

In this paper, we combine neural networks with Gene Ontology for reducing the dimensions of scRNA-seq data, which can improve the clustering of scRNA-seq data. We propose two models GOAE and GONN that are unsupervised and supervised model respectively. The proposed model mainly contains two key components: selection of significant GO terms and combination GO terms with neural networkbased model. Performance evaluation on two datasets shows that GONN and GOAE perform better than existing state-of-the-art methods.

## Author’s contributions

JP and XS designed the algorithm; XW implemented the algorithm; JP and XW wrote this manuscript. All authors read and approved the final manuscript.

## Acknowledgements

This work was supported by National Natural Science Foundation of China (No. 61702421, 61332014, 61772426), China Postdoctoral Science Foundation (No. 2017M610651), Fundamental Research Funds for the Central Universities (No. 3102018zy033).

## References

1. Wang, Z., Gerstein, M., Snyder, M.: Rna-seq: a revolutionary tool for transcriptomics. Nature reviews genetics 10(1), 57 (2009)

2. Stegle, O., Teichmann, S.A., Marioni, J.C.: Computational and analytical challenges in single-cell transcriptomics. Nature Reviews Genetics 16(3), 133 (2015)

3. Raj, A., van Oudenaarden, A.: Nature, nurture, or chance: stochastic gene expression and its consequences. Cell 135(2), 216–226 (2008)

4. Kolodziejczyk, A., Kim, J.K., Svensson, V., Marioni, J., Teichmann, S.: The technology and biology of single-cell rna sequencing. Molecular Cell 58(4), 610–620 (2015)

5. Tang, F., Barbacioru, C., Wang, Y., Nordman, E., Lee, C., Xu, N., Wang, X., Bodeau, J., Tuch, B.B., Siddiqui, A.: mrna-seq whole-transcriptome analysis of a single cell. Nature Methods 6(5), 377–382 (2009)

6. Wu, A.R., Neff, N.F., Kalisky, T., Dalerba, P., Treutlein, B., Rothenberg, M.E., Mburu, F.M., Mantalas, G.L., Sim, S., Clarke, M.F.: Quantitative assessment of single-cell rna-sequencing methods. Nature Methods 11(1) (2014)

7. Jaitin, D.A., Kenigsberg, E., Keren-Shaul, H., Elefant, N., Paul, F., Zaretsky, I., Mildner, A., Cohen, N., Jung, S., Tanay, A.: Massively parallel single-cell rna-seq for marker-free decomposition of tissues into cell types. Science 343(6172), 776–779 (2011)

8. Chung, W., Eum, H.H., Lee, H.O., Lee, K.M., Lee, H.B., Kim, K.T., Ryu, H.S., Kim, S., Lee, J.E., Park, Y.H.: Single-cell rna-seq enables comprehensive tumour and immune cell profiling in primary breast cancer. Nature Communications 8, 15081 (2017)

9. Wold, S., Esbensen, K., Geladi, P.: Principal component analysis. Chemometrics & Intelligent Laboratory Systems 2(1), 37–52 (1987)

10. Buettner, F., Natarajan, K.N., Casale, F.P., Proserpio, V., Scialdone, A., Theis, F.J., Teichmann, S.A., Marioni, J.C., Stegle, O.: Computational analysis of cell-to-cell heterogeneity in single-cell rna-sequencing data reveals hidden subpopulations of cells. Nature Biotechnology 33(2), 155–60 (2015)

11. Shalek, A.K., Satija, R., Adiconis, X., Gertner, R.S., Gaublomme, J.T., Raychowdhury, R., Schwartz, S., Yosef, N., Malboeuf, C., Lu, D.: Single-cell transcriptomics reveals bimodality in expression and splicing in immune cells. Nature 498(7453), 236 (2013)

12. Lin, C., Jain, S., Kim, H., Barjoseph, Z.: Using neural networks for reducing the dimensions of single-cell rna-seq data. Nucleic Acids Research 45(17), 156 (2017)

13. Li, X., Chen, W., Chen, Y., Zhang, X., Gu, J., Zhang, M.Q.: Network embedding-based representation learning for single cell rna-seq data. Nucleic Acids Research 45(19), 166 (2017)

14. Maaten, L., Hinton, G.: Visualizing data using t-sne. Journal of Machine Learning Research 9(2605), 2579–2605 (2008)

15. Yau, C., Pierson, E.: Dimensionality reduction for zero-inflated single cell gene expression analysis. Genome Biology 16(1), 241 (2015)

16. Xu, C., Su, Z.: Identification of cell types from single-cell transcriptomes using a novel clustering method. Bioinformatics 31(12), 1974–1980 (2015)

17. Kiselev, V.Y., Kirschner, K., Schaub, M.T., Andrews, T., Yiu, A., Chandra, T., Natarajan, K.N., Reik, W., Barahona, M., Green, A.R., et al.: Sc3: consensus clustering of single-cell rna-seq data. Nature methods 14(5), 483 (2017)

18. Ma, J., Yu, M.K., Fong, S., Ono, K., Sage, E., Demchak, B., Sharan, R., Ideker, T.: Using deep learning to model the hierarchical structure and function of a cell. Nature methods 15(4), 290 (2018)

19. Carbon, S., Ireland., A., Mungall, C.J., Shu, S.Q., Marshall, B., Lewis, S., Hub, T.A.: Amigo: online access to ontology and annotation data. Bioinformatics 25(2), 288–289 (2009)

20. Peng, J., Hui, W., Shang, X.: Measuring phenotype-phenotype similarity through the interactome. BMC bioinformatics 19(5), 114 (2018)

21. Peng, J., Zhang, X., Hui, W., Lu, J., Li, Q., Liu, S., Shang, X.: Improving the measurement of semantic similarity by combining gene ontology and co-functional network: a random walk based approach. BMC systems biology 12(2), 18 (2018)

22. Peng, J., Wang, H., Lu, J., Hui, W., Wang, Y., Shang, X.: Identifying term relations cross different gene ontology categories. BMC bioinformatics 18(16), 573 (2017)

23. Peng, J., Xue, H., Shao, Y., Shang, X., Wang, Y., Chen, J.: A novel method to measure the semantic similarity of hpo terms. International Journal of Data Mining and Bioinformatics 17(2), 173–188 (2017)

24. Peng, J., Wang, T., Wang, J., Wang, Y., Chen, J.: Extending gene ontology with gene association networks. Bioinformatics 32(8), 1185–1194 (2015)

25. Comon, P.: Independent Component Analysis, a New Concept?, pp. 287–314. Elsevier North-Holland, Inc., ??? (1994)

26. Žurauskienė, J., Yau, C.: pcareduce: hierarchical clustering of single cell transcriptional profiles. Bmc Bioinformatics 17(1), 140 (2016)

27. Vincent, P., Larochelle, H., Lajoie, I., Bengio, Y., Manzagol, P.A.: Stacked denoising autoencoders: Learning useful representations in a deep network with a local denoising criterion. Journal of Machine Learning Research 11(12), 3371–3408 (2010)

28. Pollen, A.A., Nowakowski, T.J., Shuga, J., Wang, X., Leyrat, A.A., Lui, J.H., Li, N., Szpankowski, L., Fowler, B., Chen, P.: Low-coverage single-cell mrna sequencing reveals cellular heterogeneity and activated signaling pathways in developing cerebral cortex. Nature Biotechnology 32(10), 1053–1058 (2014)

29. Sasagawa, Y., Nikaido, I., Hayashi, T., Danno, H., Uno, K.D., Imai, T., Ueda, H.R.: Quartz-seq: a highly reproducible and sensitive single-cell rna sequencing method, reveals non-genetic gene-expression heterogeneity. Genome Biology,14,4(2013-04-17) 14(4), 3097 (2013)

30. Deng, Q., Ramsköld, D., Reinius, B., Sandberg, R.: Single-cell rna-seq reveals dynamic, random monoallelic gene expression in mammalian cells. Science 343(6167), 193–196 (2014)

31. Hubert, L., Arabie, P.: Comparing partitions. Journal of Classification 2(1), 193–218 (1985)

32. Vinh, N.X., Epps, J., Bailey, J.: Information Theoretic Measures for Clusterings Comparison: Variants, Properties, Normalization and Correction for Chance, pp. 1073–1080 JMLR.org, ??? (2010)

33. Sene, K.H., Porter, C.J., Palidwor, G., Pereziratxeta, C., Muro, E.M., Campbell, P.A., Rudnicki, M.A., Andradenavarro, M.A.: Gene function in early mouse embryonic stem cell differentiation. Bmc Genomics 8(1), 85 (2007)

34. Cruz, D.SG.D., Lima, A.P.N.D., Neto, J.P., Massoco, C.: Effects of unilateral cervical vagotomy on murine dendritic cells. American Journal of Immunology (2015)

35. Pawel, K., Vijay, C., Carsten, P.: Simulating the mammalian blastocyst - molecular and mechanical interactions pattern the embryo. Plos Computational Biology 7(5), 1001128 (2011)

36. Ko, M.S.H., Zalzman, M., Sharova, L.V.: Methods for enhancing genome stability and telomere elongation in embryonic stem cells. US (2015)

